# Impact of MMP-2 and MMP-9 activation on wound healing, tumor growth and RACPP cleavage

**DOI:** 10.1101/327791

**Authors:** Dina V. Hingorani, Csilla N. Lippert, Jessica L. Crisp, Elamprakash N. Savariar, Jonathan P.C. Hasselmann, Christopher Kuo, Quyen T. Nguyen, Roger Y. Tsien, Michael A. Whitney, Lesley G. Ellies

**Author notes:** These authors contributed equally to the manuscript. Deceased August 24, 2016.

## Abstract

Matrix metalloproteinases-2 and -9 (MMP-2/-9) are key tissue remodeling enzymes that have multiple overlapping activities critical for wound healing and tumor progression *in vivo*. To overcome issues of redundancy, we created MMP-2/-9 double knockout (DKO) mice in the C57BL/6 background to examine wound healing. We then bred the DKO mice into the polyomavirus middle T (PyVmT) model of breast cancer to analyze the role of these enzymes in tumorigenesis. Breeding analyses indicated that significantly fewer DKO mice were born than predicted by Mendelian genetics and weaned DKO mice were growth compromised compared with wild type (WT) cohorts. Epithelial wound healing was dramatically delayed in adult DKO mice and when the DKO was combined with the PyVmT oncogene, we found that the biologically related process of mammary tumorigenesis was inhibited in a site-specific manner. To further examine the role of MMP-2/-9 in tumor progression, tumor cells derived from WT or DKO PyVmT transgenic tumors were grown in WT or DKO mice. Ratiometric activatable cell penetrating peptides (RACPPs) previously used to image cancer based on MMP-2/-9 activity were used to understand differences in MMP activity in WT or knockout syngeneic tumors in WT and KO animals. Analysis of an MMP-2 selective RACPP in WT or DKO mice bearing WT and DKO PyVmT tumor cells indicated that the genotype of the tumor cells was more important than the host stromal genotype in promoting MMP-2/-9 activity in the tumors in this model system. Additional complexities were revealed as the recruitment of host macrophages by the tumor cells was found to be the source of the tumor MMP-2/-9 activity and it is evident that MMP-2/-9 from both host and tumor is required for maximum signal using RACPP imaging for detection. We conclude that in the PyVmT model, the majority of MMP-2/-9 activity in mammary tumors is associated with host macrophages recruited into the tumor rather than that produced by the tumor cells themselves. Thus therapies that target tumor-associated macrophage functions have the potential to slow tumor progression.

## Introduction

Tissue matrix homeostasis is a complex process that is important in normal growth, development and wound healing. Matrix metalloproteinases-2 and -9 (MMP-2/-9) are members of a family of over 25 zinc-dependent endopeptidases that degrade or cleave a wide range of extracellular proteins including components of the extracellular matrix (ECM). Proteolysis is regulated at multiple levels, including transcription, secretion, and conversion of the zymogen (pro-MMP) into an active protease as well as by the presence of cell type specific tissue inhibitors of metalloproteinases (TIMPs) (1, 2). Elevated MMP-2/-9 levels are associated with proinflammatory states that can induce or amplify diseases, such as cardiac disease, arthritis and cancer (3–5), suggesting a role for inhibitors in disease prevention or treatment.

Early efforts to develop therapeutic inhibitors were met with disappointment. This was due to side effects from insufficiently specific inhibitors as well as an inadequate understanding of the normal functions of these enzymes and the complex interactions taking place *in vivo* (6, 7). Evidence now suggests that MMPs act as key nodal components of an interconnected protease web and they can have opposing effects on the same biological process depending on factors present in the local microenvironment (8). For example, it is now recognized that many MMPs, including MMP-2/-9, can be protective in cancer and that their upregulation may be involved in processes aimed at eliminating abnormal tumor cells. Regardless of the function of MMPs in cancer, fluorescence activatable probes that rely on MMP activity have been developed to visualize tumor margins and improve surgical outcomes (9–11).

A number of different genetically engineered mouse models have been used to improve our understanding of the complex interactions occurring between MMPs and their *in vivo* microenvironments (8, 12, 13). Because MMP-2/-9 have overlapping functions *in vivo*, we used double mutant mice to study the role of these enzymes in wound healing and tumorigenesis. We also used imaging probes dependent on MMP-2/-9 activity to identify cell types within tumors where the activity was greatest. Our findings reveal that tumor cells play a critical role in recruiting host stromal cells that activate MMP-2/-9 *in vivo* in our model system.

## Materials and Methods

### Mice

We backcrossed both the MMP2^−/−^ (14) and MMP9^−/−^ mice (15) (a generous gift from Lisa Coussens) until they were congenic on an albino C57Bl/6 background. The mice were then mated to produce MMP-2/-9 double knockout (DKO) mice. Because DKO matings were not fertile, we bred one DKO with an MMP2^+/−^MMP9^−/−^ mate. The DKO and heterozygous/KO mice could be of either sex in the breeding pair. Wild type (WT) albino C57Bl/6 mice were used as controls for the DKO strain since WT littermates were not generated in these complex breedings. To examine mammary tumorigenesis, DKO mice were bred into the polyomavirus middle T (PyVmT) model of mammary tumorigenesis [B6.FVB-Tg(MMTV-PyVT)634Mul/LellJ; The Jackson Laboratory, Bar Harbor, ME] (16) on an albino C57Bl/6 background. When tumor-bearing animals were euthanized, the tumors and mammary fat pads were excised and weighed. The mammary fat pads were formalin fixed and stained with carmine as previously described (16). All animal studies were performed in compliance with the recommendations in the Guide for the Care and Use of Laboratory Animals of the National Institutes of Health. The protocols were approved by the Institutional Animal Care and Use Committee of UC San Diego (Protocol numbers: S01162, S04011). All surgery was performed under isoflurane anesthesia and all efforts were made to minimize suffering.

### Wound healing

Bilateral 8 mm full thickness skin incisions were made on the dorsal surface of the flank on either side of the spine in 6 mice per group. The wound was sutured closed with resorbable sutures. On day 11, the superficial wound area, including any unhealed scab region, was measured and the mice were euthanized. The skin was fixed in formalin and paraffin embedded; then, crosssections along the initial wound line at approximately the same vertical location were stained with hematoxylin and eosin (H&E). As an additional measure of wound healing, the distance between healthy hair follicles on the cross sections was quantified.

### Real time PCR

Total RNA was isolated from mammary tumors using the RNeasy kit (Qiagen, Valencia, CA) according to the manufacturer’s instructions. First-strand cDNA was synthesized using an MMLV-Reverse Transcriptase and random hexamers (Promega, Madison, MI) and then amplified using primers listed in Table S1. Semi-quantitative real time PCR was performed using EvaGreen SYBR Green mastermix (Fisher Scientific, Houston, TX) in a Step One Plus Real time PCR system (Applied Biosystems, Grand Island, NY). The gene expression levels were calculated after normalization to the standard housekeeping genes, ß actin and glyceraldehyde 3 phosphate dehydrogenase (GAPDH), using the ΔΔ*C*_T_ method and expressed as the relative mRNA level compared with the internal control.

### PyVmT cell lines

Cell lines were derived from PyVmT WT or DKO transgenic tumors in the C57Bl/6 background as previously described (17, 18). Tumor cells (10^5^ to 10^6^) were orthotopically injected into the pectoral mammary glands of adult female mice in 2 mg/ml Matrigel (BD Biosciences, San Jose, CA). Because the individual cell lines had different growth rates, we injected the cell lines at different times in an attempt to make the tumors a uniform size for imaging.

### RACPP cleavage

RACPPs were synthesized using standard solid phase 9-fluorenylmethyloxycarbonyl (Fmoc) synthesis and characterized as previously reported (19). Briefly, a fluorescent donor and acceptor are placed onto polycationic and polyanionic domains, respectively, in sufficient proximity for fluorescence resonance energy transfer (FRET). If the linker between the polycation and polyanion is cut, typically by a protease, the two halves of the RACPP dissociate, immediately causing disruption of FRET and a large increase in the ratio of donor to acceptor emissions. Also the polycation is taken up and retained at or near the site of proteolysis, while the polyanion is subject to pharmacokinetic washout, reinforcing the high ratio of donor to acceptor emissions. As described in (19) the cleavable RACPP, with a PLGC(Me)AG linker sequence and referred to as RACPP1 in the cited paper, could be cut by MMP2 (Kcat/Km=36429 s^−1^M^−1^), MMP-9 (Kcat/Km=13503 s^−1^M^−1^), and somewhat by MMP14 (Kcat/Km=17173 s^−1^M^−1^), whereas the uncleavable control RACPP was resistant to such cleavage (19, 20). We also synthesized an RACPP with a cleavable sequence TLSLEH in the manner described above. This sequence is selective for MMP2 (Kcat/Km=11405 s^−1^M^−1^) and slightly cleaved by MMP14 (Kcat/Km=1200 s^−1^M^−1^) uncleaved by the related gelatinase MMP9. WT and DKO breast cancer cells (10^5^ to 10^6^ in 2 mg/ml Matrigel) were orthotopically injected into the mammary fat pads of albino C57BL/6 WT and DKO mice. When both the WT and DKO tumors were palpable, 10nmol of RACPPs was dissolved in 100μl sterile water (Conc=100uM) and administered intravenously (retroorbital) while mice were under isoflurane anesthesia. Two hours after peptide administration, mice were euthanized by isoflurane overdose and then cervical dislocation. The skin was removed and mice were imaged as previously described (19). Briefly, Cy5 was excited at 620/20 nm and the emission intensity was measured in 10-nm increments, ranging from 640-680 nm, through a tunable crystal emission filter. Numerator (Cy5) and denominator (Cy7) values were generated by integrating the spectral images over 660-720 nm and 760-830 nm, respectively. Custom software divided the Cy5 emission by the Cy7 emission to create a pseudocolor ratio value image, ranging from blue (lowest ratio) to red (highest ratio). Ratios were quantitated using ImageJ. To compare the data from two independent sets of experiments in albino C57BL/6 mice, the ratios were normalized, adjusting the values for each separate experiment by dividing each ratio value by the lowest ratio for that experiment of mice (as a result, the lowest ratio for each experiment was set to one).

### Immunofluorescence

Perfusion fixed py8119-lentiGFP tumor samples from mice that were treated with Cy5:Cy7 RACPP were suspended in 20% sucrose solution overnight at 4C prior to embedding in OCT solution. 10μm sections were made and treated with a 1:1000 dilution of Alexa405 conjugated primary antibody to F4/80 marker for macrophages (Abcam, Cambridge, UK). The slides were placed in a humififier chamber overnight at 4C followed by washes with PBS and coverslipped. Three color confocal imaging was performed using the Nikon A1 system with laser lines 405nm, 488nm and 640nm.

### Immunohistochemistry

To further determine whether the tumor cells or host stroma contributed more to the MMP-2/-9-cleavable RACPP ratios, we cryosectioned (10 μm) and then imaged the tumors harvested from the C57Bl/6 experiment using a confocal microscope (Nikon Instruments Inc, Melville, NY). Additional sections (5 μm) were stained for neutrophils using the NIMP-R14 antibody (Abcam, Cambridge, UK) with standard immunohistochemical (IHC) methods. Since macrophages are a major source of MMP-2/-9 activity, we examined their infiltration into tumors. Formalin fixed, paraffin embedded tumor samples were sectioned at 7 μm and stained with the F4/80 antibody (BM8; eBioscience, San Diego, CA, dilution 1:200) following antigen retrieval in citrate buffer pH 6.0, 0.05% tween 20. Antibody visualization was with ImmPACT DAB staining (Vector Laboratories Inc, Burlingame, CA). Slides were scanned using a Nanozoomer and analyzed using Aperio Imagescope software (Leica Biosystems Inc, Buffalo Grove, IL).

### Statistics

Breeding results were analyzed using the Chi-square test. Normally distributed data were analyzed using the Student’s t test or by ANOVA followed by multiple comparisons using the Holm-Sidak correction. They are presented as means ± SEM. Nonparametric data were analyzed using the Mann-Whitney test. All data were analyzed using Graphpad Prism software (Graphpad Prism, La Jolla, CA).

## Results and Discussion

### Reduced fecundity and compromised growth in DKO mice

Given the fundamental roles that MMP-2/-9 play in tissue homeostasis, it is reasonable to hypothesize that a loss of both enzymes could result in reduced fertility and offspring viability. Furthermore, while it has been shown that MMP-2 deficiency does not affect breeding success (14), the loss of MMP-9 results in smaller litter sizes and an increased percentage of infertile breeding pairs (21). However, these changes do not appear to be due to impaired embryonic and fetal development as heterozygous matings resulted in the expected Mendelian frequencies of MMP9^+/+^, MMP9^+/−^ and MMP9^−/−^ mice (15, 22). Similarly, we observed a reduced litter size in our DKO mice as follows: WT breeding 6.27 pups ± 0.31 vs DKO breeding 4.76 pups ± 0.31 (mean ± SEM, p < 0.001). However, only 70% of the DKO mice that were expected according to Mendelian ratios survived to weaning (Table S1). These results indicate a significant functional overlap between the two enzymes in reproduction such that the DKO exacerbates the MMP-9 null phenotype, skewing the normal Mendelian ratios and reducing the number of viable DKO mice.

Although no significant difference in survival of weaned male and female DKO mice was observed, we found mild but significant early growth retardation in DKOs of both sexes (Figure S1), which has not been observed in single KO mice. Male DKO mice were more compromised at an early age compared with their WT counterparts (~57% reduction in body weight at 3-6 weeks, recovering to 86% by 12 weeks; Figure S1A) than female DKO mice (~84% reduction in body weight from week 6; Figure S1B), underscoring the important role MMP-2/-9 play in normal development.

### DKO mice have delayed wound healing

At a cellular level, wound healing has much in common with normal development, and numerous studies suggest that MMP-2/-9 play active roles in this process (23). Accordingly, we observed a delay in primary wound healing in DKO mice compared with WT mice. First, at 11 days post-incision, wound areas for the DKO mice were significantly larger (9.67 ± 2.09 mm^2^) than those for the WT mice (0.12 ± 0.03 mm^2^; mean ± SEM, p <0.001; n = 6 mice [total of 12 wounds] per group) (Figure 1A-B). Additionally, even though two vertical incisions were made to create the wound, the scab area that formed as part of the wound healing process crossed from one incision side to the other for some DKO mice (Figure 1A), while the WT mice were almost completely healed by Day 11. This may be attributed to a number of factors that occur during the healing period. First, although MMP expression in healthy skin is low (24), MMP-9 expression can be induced at the leading edge of migrating epithelial cells, enabling these cells to move through the ECM and re-epithelialize the wounded area (25–28). MMP expression can also be upregulated in inflammatory cells, such as macrophages, T cells and eosinophils, which infiltrate the wound and assist with pathogen clearance (29, 30). There is a complex pattern of expression involving high MMP-9 expression in the early inflammatory phase and a later increase in MMP-2 expression that occurs during the proliferative phase of wound repair (31). MMP-2 is also found in immune cells and it promotes functional recovery after spinal cord injury (32). Consistent with previous work, delayed wound healing in DKO compared to WT mice in our study was associated with aberrant re-epithelialization of the injured area, however further study is needed to clarify the overlapping mechanistic roles of MMP-2/-9 in wound healing.

**Figure 1.**
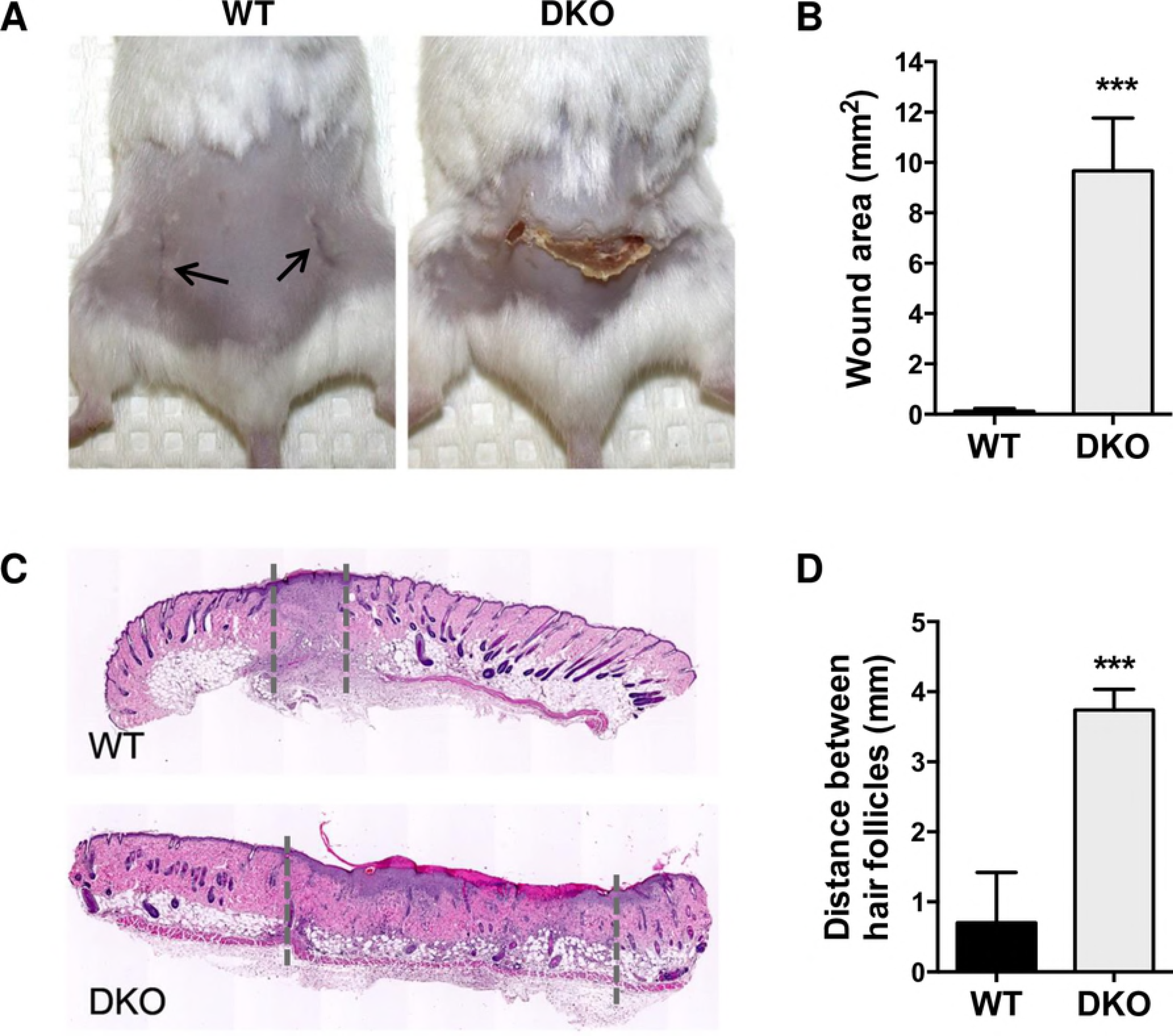
Delayed wound healing in DKO mice. A. Representative WT and DKO mice on Day 11 following the creation of bilateral, vertical, 8-mm wounds. The initial wounds were located as indicated by the arrows in the WT panel, with aberrant healing in the DKO apparent. B. Quantitation of the day 11 skin surface wound area (n = 6 mice and 12 wounds per group). C. H&E stained cross-sections of skin from WT and DKO mice at day 11 after wounding with wound margins indicated by dashed lines. D. Quantitation of the distance between healthy hair follicles adjacent to the wound (n = 2 ‒ 3 sections per wound; 12 wounds per group). Data are means ± SEM, analyzed by Student’s t test. *** p < 0.001.

To evaluate wound healing at the microscopic level, we measured the distance between healthy hair follicles. The distance between hair follicles was significantly larger for the DKO mice than for the WT mice (3.74 ± 0.3 mm v. 0.70 ± 0.13 mm; mean ± SEM, p <0.001; n = 12 wounds/group) (Figure 1C-D), indicating that wound healing had not progressed normally for the DKO mice. Interestingly, wound healing in a model of laser-induced choroidal neovascularization, mimicking human age-related macular degeneration, is nearly completely prevented in DKO mice, while the wound healing in single KOs is only partially impaired (33). This is thought to be due to an effect of MMP-2/-9 on fibrinolysis, which supports angiogenesis. Thus, inadequate vascularization may also play a role in our observation of impaired wound healing.

### Site-specific effect on tumor growth

Since MMP-2/-9 have long been associated with cancer progression either through their effects on matrix degradation or as regulators of growth factor and cytokine bioactivity (34), we next examined their role in tumor growth in the PyVmT transgenic mouse model of breast cancer (16, 35). Due to the complex breeding and relatively poor breeding success in generating PyVmT positive DKO female mice, we had a limited number of mice in this study. We used semi-quantitative real time PCR to confirm the loss of MMP-2/-9 expression. Our analysis verified that the only enzymes that were significantly downregulated in our panel of 20 MMPs were MMP-2 and MMP-9, although there was an interesting trend towards downregulation of a number of other MMPs in the absence of active MMP-2/-9. Additionally, we were unable to identify RNA from any alternative MMPs that were upregulated to compensate for the loss of MMP-2/-9 in the DKO mice compared to WT mice (Figure S2).

While no significant difference in the overall tumor burden between the PyVmT;WT and PyVmT;DKO mice was found (Figure 2A), our analysis showed a slight, but significant, reduction in tumor growth in the #4 mammary fat pad (0.431 ± 0.074 g v. 0.136 ± 0.051 g; mean ± SEM, p < 0.05) (Figure 2B). This difference could be visualized in whole mounts of the #4 mammary gland at 24 weeks of age, which showed a reduction in the amount of carmine stained mammary epithelial tissue in the PyVmT;DKO gland (Figure 2C). Interestingly, iNOS−/− mice show a similar site-specific reduction in mammary tumor growth in the inguinal fat pads (16), which are the largest of the mammary fat pads and contain a central lymph node. Since MMP-2/-9 and iNOS are key effectors of macrophages, we further investigated whether this growth retardation was primarily due to loss of MMP activity in the tumor cells or stromal cells particularly macrophages, using an orthotopic tumor cell injection model.

**Figure 2.**
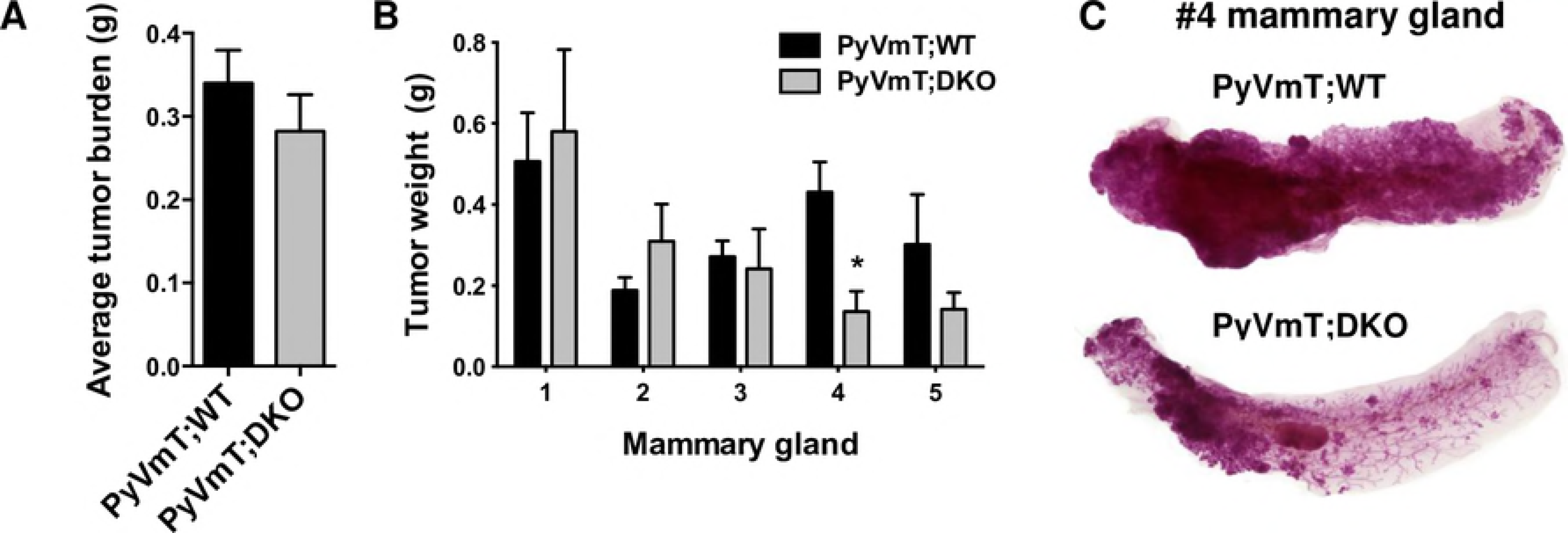
Modest effect of DKO on mammary tumorigenesis. A. Comparison of tumor burden in PyVmT;WT (N = 10) and PyVmT;DKO mice (N = 5) aged 22-25 weeks. B. Tumor burden by mammary gland site from pectoral (#1, 2, 3) to inguinal (#4, 5). Data are means ± SEM analyzed by Student’s t test, * p < 0.05. C. Whole mounts of the #4 inguinal mammary glands indicate delayed tumorigenesis in the DKO.

The results of orthotopic tumor cell injections supported a role for MMP-2/-9 in tumor growth. In one set of C57BL/6 mice used for imaging (Figure 3A), we recorded the tumor weights and observed the highest tumor weight for the WT tumor in WT mice (0.355 ± 0.09 g; mean ± SEM, p < 0.05 compared to WT tumors in DKO mice, 0.07 ± 0.06 g, DKO tumors in WT mice, 0.07 ± 0.03 g and DKO tumors in DKO mice, 0.07 ± 0.02 g) (Figure 3C). The WT tumor cell-stroma combination resulted in tumors that were almost 5-fold larger than any of the other combinations: WT tumors in DKO mice; DKO tumors in WT mice; and DKO tumors in DKO mice, suggesting that loss of MMP-2/-9 activity in either the tumor cells or the host stroma could reduce tumor growth. Based on these data, we posited that both the tumor and host stroma require MMP-2/-9 to promote tumor growth, resulting in a larger tumor size. It merits noting that in other experiments, the WT tumors were injected after the DKO tumors were already palpable. This was because the WT tumor cell line had a faster growth rate *in vitro* and *in vivo* than the DKO tumor cells, which is consistent with a tumor cell growth-promoting role for MMP-2/-9 in this strain.

**Figure 3.**
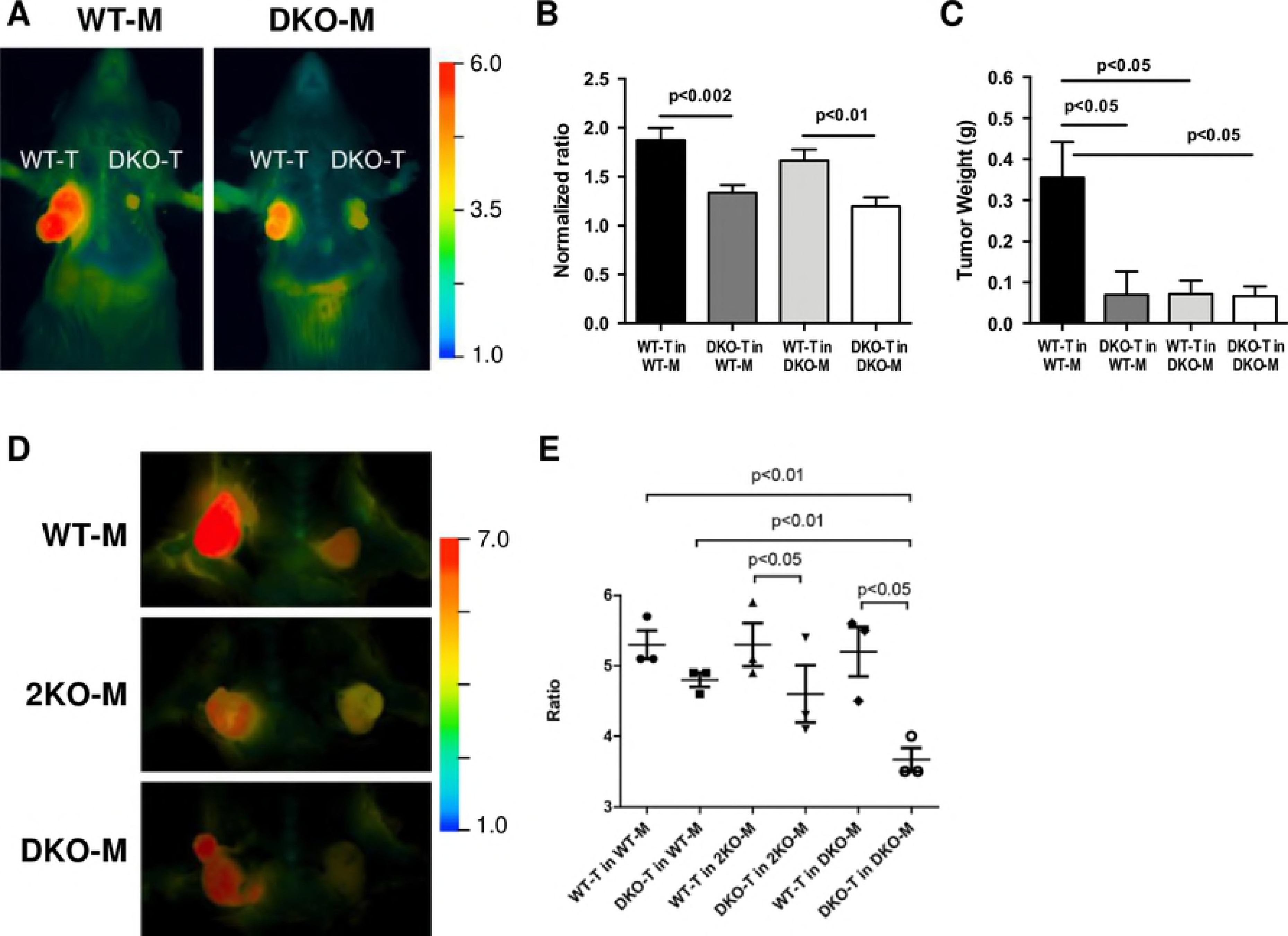
The tumor cell genotype contributes more than the stromal genotype to MMP-2/-9 activity in cleaving the MMP-2/-9-cleavable RACPP. A. C57BL/6 mice (WT and DKO) with orthotopic WT and DKO tumors (T) in their bilateral mammary fat pads. After 2 h incubation with an intravenously administered MMP-2/-9-cleavable RACPP, the tumors were imaged. B. The tumor ratios (Cy5 emission/Cy 7 emission, corresponding to cleaved/uncleaved ACPP) were quantified (N = 5 mice/group; N = 7 tumors/group). The data from two sets of independent experiments, which had the same relative comparison between tumor groups with different overall ratio ranges, were normalized so the sets could be combined. Each set was normalized to its lowest ratio (all values divided by the lowest ratio value) such that the lowest ratio for each set was re-mapped onto the value one. C. The tumor weights from one of the C57BL/6 mouse strain experiments; the weight was highest for the WT-tumor (T) in WT-mouse (M). Data are means ± SEM analyzed by one-way ANOVA and Holm-Sidak’s multiple comparisons test. D. Ratiometric images Cy5/Cy7 2 h after intravenous injection of MMP-2 selective RACPP, with cleavable sequence TLSLEH, in WT, 2KO (N = 4 mice per WT or KO group; N = 8 tumors/group) and DKO mice (N = 3 mice/group; N = 6 tumors/DKO group). In each mouse, the WT tumor is on the left and the DKO tumor is on the right. E. Quantified tumor ratios of Cy5 emission/Cy7 emission for the cohort of 11 mice imaged with the MMP2-selective RACPP, stratified by tumor type and mouse strain.

### MMP-2/-9 tumor cell genotypes contribute more than stromal genotypes to RACPP cleavage

ACPPs and, more recently, RACPPs, have proven useful in detecting protease activity in cancer (9, 11, 36, 37). A role for stromal-derived MMPs in cancer progression has become increasingly apparent as MMP expression is frequently higher in stromal cells than tumor cells (38) and MMP-2/-9 expression can be increased in stromal cells by paracrine stimulation or direct contact with malignant tumor epithelium (39). We applied this technology to our tumor cell injection model in a 4-way comparison (WT tumor cells in WT mice; DKO tumor cells in WT mice; WT tumor cells in DKO mice; and DKO tumor cells in DKO mice) to study the contribution of MMP-2/-9 stromal versus tumor cell activity in more detail (Figure 3A). The normalized Cy5/Cy7 ratio for the WT tumors in WT mice (1.87 ± 0.11; 5 mice with 7 tumors/group) was significantly higher than the ratio for the DKO tumors in WT mice (1.34 ± 0.07; p <0.003) or the DKO tumors in DKO mice (1.20 ± 0.08; p < 0.0002). The WT tumors in DKO mice (1.67 ± 0.11) also had significantly higher ratios than the DKO tumors in DKO mice (p < 0.008). The difference in ratios between the DKO tumors in WT mice and WT tumors in DKO mice did not quite reach significance (p<0.06). The WT tumors in WT mice did not have significantly higher ratios than the WT tumors in DKO mice (p = 0.25), nor did the DKO tumors in WT mice have significantly higher ratios than the DKO tumors in DKO mice (p = 0.31) (Figure 3B), suggesting that a tumor cell’s ability to activate MMP-2/-9 is more important than the host genotype to the imaging ratio. Measurement of the tumor weights indicated that MMP-2/-9 play an important role in tumor growth (Figure 3C), which is not surprising given their role in cellular migration and angiogenesis (40).

Since the DKO tumors grew at a slower rate than the WT tumors, we carried out experiments in which we injected the WT tumor cells when the DKO tumor was just palpable. We tested an MMP-2 selective RACPP (cleavable sequence TLSLEH) in WT, 2KO and DKO mice and found that loss of MMP-2 in the host significantly reduced cleavage of the probe, validating the selectivity of this cleavage sequence (Figure 3D,E). WT tumors, regardless of mouse genotype, showed high Cy5/Cy7 ratios owing to cleavage of the MMP-2 selective sequence; the WT groups and their ratios were: WT tumor in WT mice (5.3 ± 0.35); WT tumor in 2KO mice (5.3 ± 0.53) and WT tumor in DKO mice (5.2 ± 0.61). These data support the involvement of tumor derived MMP-2 in promoting high cleavage of the MMP-2 selective RACPP. However, a comparison of WT tumor (5.3 ± 0.35) and DKO tumor (4.8 ± 0.17) ratios in WT mice show statistically significant difference when either ratio is compared with the Cy5/Cy7 ratio in DKO tumor in DKO mice (3.6 ± 0.29, p<0.01). These data indicate there is a contribution from host MMPs. Overall, our data suggest that the tumor cell genotype contributes more than the stromal genotype to the detection ratios we observed for the MMP-2/-9-cleavable RACPPs in tumors.

### Host stroma influences the tumor ratio at a microscopic level

The importance of MMP-2/-9 in the tumor cells was more apparent at the microscopic level, as WT tumor cells implanted in WT mice had higher ratios than those implanted in DKO mice (4.35 ± 0.25 and 3.34 ± 0.24, respectively, p = 0.01), indicating that more subtle differences could be detected with higher magnification (Figure 4A,B). In contrast, the ratios for DKO tumors in either WT or DKO mice were not significantly different (2.13 ± 0.12 and 2.16 ± 0.13, respectively; p = 0.87). MMP-2/-9 expression in mammary carcinoma cells is associated with epithelial to mesenchymal transition and increased tumor cell invasivesness (41), so it is not surprising that our invasive WT cell line affects the RACPP ratios more than the DKO cell line. Importantly, our RACPP findings were consistent at both the macroscopic and microscopic levels. MMP-2/-9 activatable RACPPs coupled with chemotherapeutic agents have been shown to be effective in reducing breast cancer burden in animal models (42). Our results suggest this efficacy is due in part to the ability of these agents to target both tumor cells and their associated tumor-promoting stroma.

In addition to examining the ratios at a microscopic level, we examined the tumor morphology after H&E staining and found that while the WT tumors were dense with tumor cells, the DKO tumors had a looser tissue organization (Figure 4A). The differences in growth rates, which reflect tumor heterogeneity and different mammary tumor subtypes explains the differences in their morphology (18).

**Figure 4.**
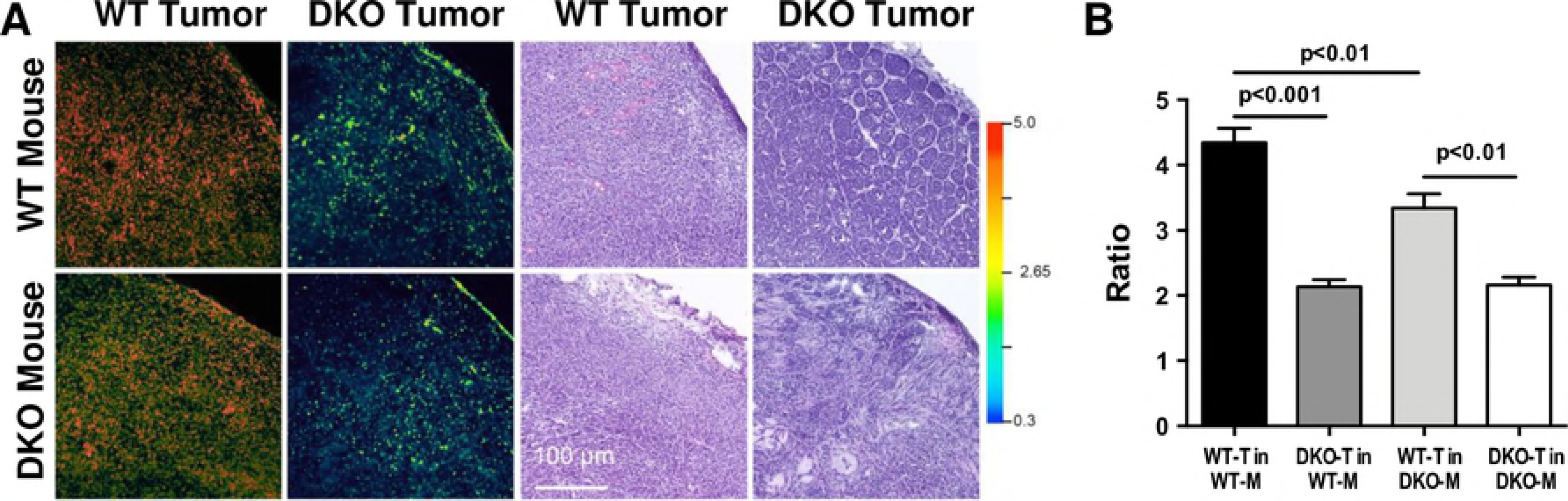
At the microscopic level, the host stroma enhances the WT-T ratio. A. The ratios for tumor sections (10 μm; WT-T in WT-M and DKO-M and DKO-T in WT-M and DKO-M) were evaluated with confocal microscopy and then the tissue sections were stained with H&E to examine the morphology. B. Quantitation of the ratios corresponding to 4 ‒ 6 confocal images per group. Data are means ± SEM analyzed by one-way ANOVA and Sidak’s multiple comparisons test.

Since the infiltrating cells had the morphology of macrophages, we carried out experiments in which Py8119GFP tumors were injected with RACPPs and later stained with the macrophage marker F4/80 (Figure 5A). We found that indeed, the cells with high RACPP signal at the periphery of the tumors were macrophages (Figure G,H). Macrophages were distributed throughout the stromal areas surrounding the tumor cells and showed accumulation of RACPP (Figure 5B-D). RACPP positive macrophages present in the center of the tumors showed less cleavage, likely due to reduced MMP-2/-9 activity (Figure 5I-J). At higher magnification, we were able to clearly show that the tumor cells were also positive for RACPP, but at a lower intensity than the macrophages (Figure 5E-L). However, because of the abundance of the tumor cells, they contribute significantly to the total RACPP signal observed (Figure E,F). Our data indicate that RACPPs are useful tools to localize MMP-2/-9 activity in vivo and confirm that MMP-2/-9 activity is present in both tumor cells and macrophages.

**Figure 5.**
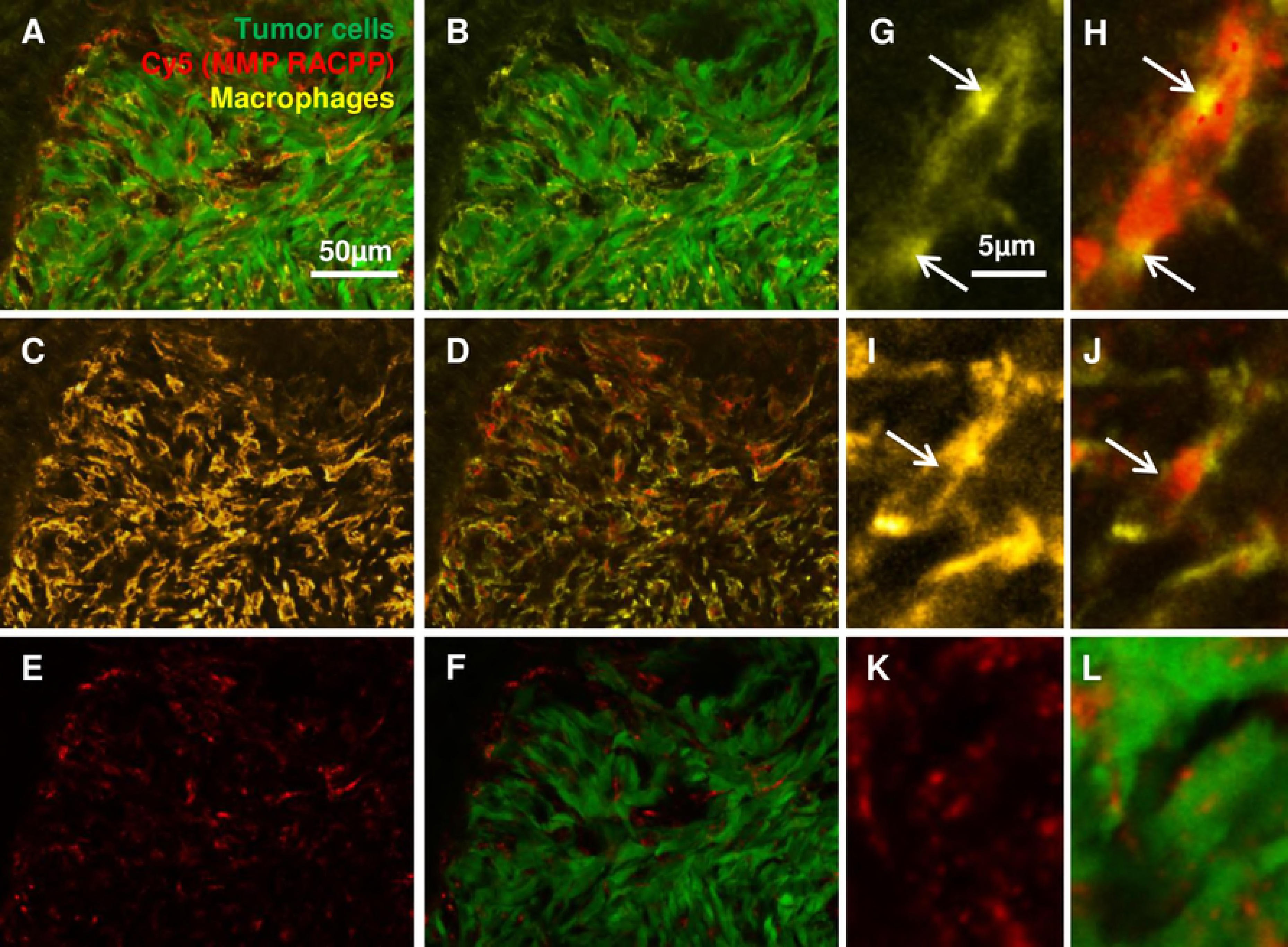
RACPP ratios are higher in macrophages than tumor cells. A. Immunofluorescence staining with F4/80 pan macrophage antibody marker (yellow) on Py8119-GFP (green) tumor tissue excised from C57Bl6-albino mice injected with MMP cleavable RACPP (Cy5: red). B. Macrophage infiltration surrounding tumor cells. C. Macrophage distribution in the tissue and D. overlay with Cy5 from cleaved RACPP due to MMP-2/-9 activity. E,F. Much of the Cy5 from cleaved RACPP is seen in stromal region surrounding tumor cells with dimmer puncta seen on the Cy5 image alone. G,H. Higher magnification images demonstrating high Cy5 signal in the macrophages at the tumor periphery rather than I, J. those at the tumor center. K,L. Higher magnification showing accumulation of Cy5 from cleaved RACPP in the tumor cells.

To further understand the ratio enhancement for WT tumors in WT mice (v. DKO mice), we quantitated the number of macrophages infiltrating the tumors. As expected, the majority of the infiltration was at the periphery of the tumor and WT tumor cells had a significantly greater ability to recruit host macrophages into the tumor than DKO tumor cells (Figure 6A-F). Interestingly, a lack of MMP-2 or MMP-2/-9 in the host tissues reduced the effectiveness of WT tumor cells in recruiting macrophages (Figure 6G), which is possibly due to a motility defect in the KO macrophages. DKO mice also failed to recruit macrophages in the choroidal neovascularization model although the mechanism was not determined (33). The effects of the KO mice on macrophage recruitment into WT tumors was less prominent in the tumor center (Figure 6H), possibly due to the presence of a subpopulation of macrophages that were not dependent on MMP-2/-9 for motility. Overall, our results in mouse mammary tumors are consistent with previous studies showing that human colorectal cancer cells induce stromal macrophage MMP-2/-9 production (43, 44).

**Figure 6.**
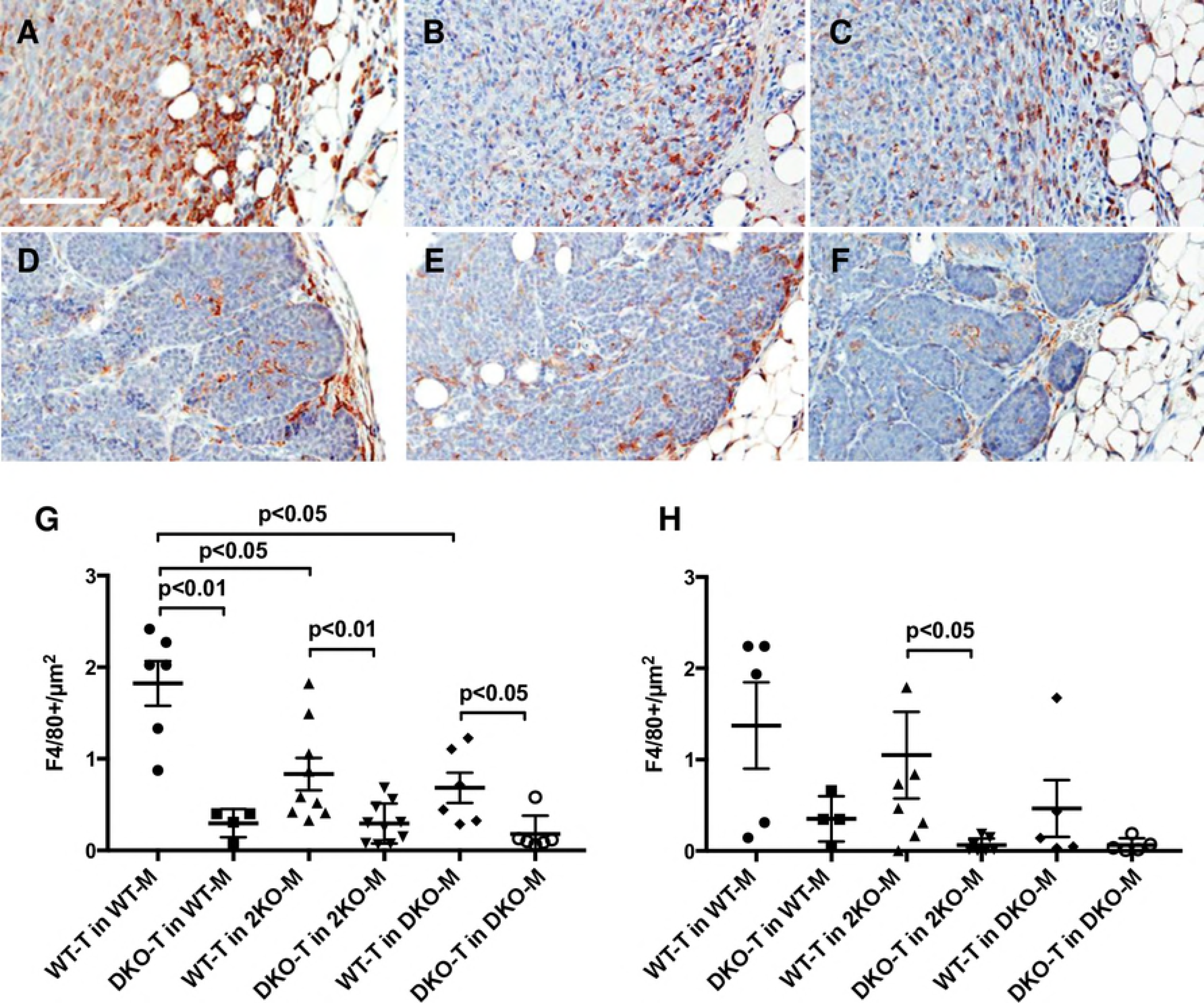
Tumor associated macrophage (TAM) infiltration is modulated by MMP-2/-9. Panels A-F are representative sections stained with F4/80 to identify TAM infiltration in the various tumor cell/host mouse genotype combinations. A. WT-T in WT mouse. B. WT-T in 2KO mouse. C. WT-T in DKO mouse. D. DKO-T in WT mouse. E. DKO-T in 2KO mouse. F. DKO-T in DKO mouse. Scale bar 100 μm. G. Macrophage infiltration at the periphery (0.5 mm into the tumor). H. Macrophage infiltration into the center of the tumor (1 mm in from the tumor boundary). N = 6-8 tumors per condition. Data are means ± SEM.

We also evaluated the number of neutrophils, which produce MMP-2/-9, in tumor sections (Figure S3). Neutrophils were a potential candidate cell type contributing to additional activity in the WT tumor in WT mice because neutrophils secrete MMP-9 without tissue inhibitor metalloproteinase 1 (TIMP-1), whereas many other cell types secrete MMP-9 and TIMP-1 together(45). Neutrophil counts had a trend towards higher levels in WT mice than DKO mice (16.21 ± 4.05 v. 8.25 ± 2.06 neutrophils / mm^2^, p <0.07), but it was clear that macrophages were the major myeloid cells present in the tumors.

## Conclusions

Our data comparing WT and DKO mice confirm an important role for MMP-2/-9 in wound healing and tumorigenesis. In the *in vivo* tumor microenvironment, our data show that the majority of MMP-2/-9 activity is associated with the tumor cell genotype in our model system. The complex interplay between tumor cells and host cells is exemplified by the data indicating that WT host macrophages recruited by WT tumor cells play an important role in RACPP cleavage. While a new generation of more specific MMP-2/-9 inhibitors has been developed and is undergoing clinical trials (7), several alternative strategies targeting macrophage activity have been proposed. Exploiting drugs that inhibit macrophage recruitment into tumors (46), harnessing macrophage Fc *γ* R-mediated processing for local delivery of antibody-drug conjugates (47), or macrophage mediated drug delivery to the tumor’s extracellular matrix (47) may prove beneficial in slowing tumor progression. RACPP-drug conjugates can be selectively delivered to tumors (42, 48) and our results confirm that they can be processed both in tumor cells and tumor-associated macrophages to provide therapeutic benefits.

## Acknowledgments

We thank Kathryn Talisman for animal husbandry and Paul Steinbach for assistance with imaging. This work was supported by NIH grants K22CA118182 (LGE), R01CA158448 (RYT), and P30NS047101 (UCSD Microscopy Core).

## Supporting Information

**Table S1.**Breeding results for production of MMP-2 and -9 double KO mice. * p < 0.05 Chi-square test. The number of pups reflects those surviving to weaning at 3 weeks of age.

**Figure S1. Growth is compromised in DKO mice.** A. Average body weights of wild type (WT) and double KO (DKO) mice. B. Average body weights of WT and DKO female mice with individual body weights of DKO females on the right. Data are means ± SEM analyzed by t tests using the Holm-Sidak correction for multiple comparisons. N = 7-14 mice per group. * p<0.001

**Figure S2. MMP-2 and -9 deletion was confirmed by RT-PCR.** Semi-quantitative RT-PCR showing the gene expression of a panel of MMPs. MMP-2/-9 are the only MMPs with significant differences between the WT and DKO tumors. N = 6 tumors from PyVmT;WT or PyVmT;DKO mice. Data are box and whisker plots with min and max, ** p<0.01, Mann-Whitney test.

**Table S2. Oligonucleotides**

**Figure S3. Neutrophil infiltration in WT and DKO tumors.** A. Immunohistochemistry for neutrophil staining (NIMP-R14 antibody) of tumor sections with no primary antibody (control) as well as in WT-T in WT-M and DKO-M. B. The brown staining neutrophils from immunostained tumor sections were counted and represented per mm2 of tumor area (4-6 sections/group). Data are means ± SEM analyzed by Student’s t test.

